# Atomic Protein Structure Modeling from Cryo-EM Using Multi-Modal Deep Learning and AlphaFold3

**DOI:** 10.1101/2025.03.16.643561

**Authors:** Nabin Giri, Jianlin Cheng

## Abstract

Cryo-electron microscopy (cryo-EM) has revolutionized structural biology by enabling near-atomic resolution visualization of protein structures. However, accurately modeling 3D atomic structures from cryo-EM density maps remains challenging, particularly for large multi-chain complexes and assemblies. Here, we present an automated pipeline that integrates multi-modal deep learning and advanced structure prediction techniques to improve model accuracy. Our approach leverages a deep learning model that combines sequence-based features from a Protein Language Model with cryo-EM density maps, enabling a richer feature representation across modalities. The deep learning-predicted voxels are utilized to build a Hidden Markov Model (HMM) and a tailored Viterbi algorithm is used to align sequences to generate an initial protein backbone structures. These backbone models then serve as templates for AlphaFold3, which refines the structures for improved accuracy. Our approach combines cryo-EM data with AlphaFold3 predictions, helping to refine and improve AlphaFold3’s predicted structures. By integrating both methods, we can generate more accurate and reliable atomic models, particularly for large proteins with complex conformations.

## Introduction

Determining the three-dimensional (3D) atomic structures of macromolecules, such as protein complexes and assemblies, is important for understanding biological functions at the molecular level. The spatial arrangement of atoms within these macromolecules provides key insights into protein structure, function, and interactions, guiding research in structural biology and rational drug discovery [1, 2, 3, 4, 5]. In recent years, cryo-electron microscopy (cryo-EM) has emerged as a powerful experimental technique for visualizing large protein structures at near-atomic resolution. However, accurately building atomic structures from high-resolution cryo-EM density maps remains a challenging task due to missing electron values and local low resolution in some regions of cryo-EM maps which makes interpretation difficult.

Several computational approaches, including Phenix [6], DeepTracer [7], DeepMainmast [8], Cryo2Struct [9], and ModelAngelo [10], provide automated solutions for modeling atomic structures within cryo-EM density maps [11, 12]. Among them, Phenix [6] is a widely used tool that utilizes classical molecular optimization techniques to build atomic protein structures. DeepTracer [7] provides a deep learning-based web platform for automated structure building, while ModelAngelo [10] integrates cryo-EM data, amino acid sequences, and prior knowledge of protein geometries to refine structural models and assign residue identities. More recently, DeepMainmast [8] has combined AlphaFold2 [13] predictions with density-guided tracing protocols, significantly improving atomic model building from cryo-EM maps. These advancements have demonstrated that incorporating AlphaFold-predicted structures improves the reliability and accuracy of cryo-EM-based atomic structure determination.

Building on this momentum, we introduce a versatile workflow that integrates cryo-EM density data with AlphaFold3 [14] predictions, further refining and improving the accuracy of AlphaFold3-generated structures. By leveraging cryo-EM templates in the AlphaFold3 framework, our approach allows the generation of more accurate atomic models, particularly for proteins with flexible conformations and regions of low-resolution density. This integration improves the structure prediction capabilities, facilitating the modeling of complex biomolecular assemblies and expanding the potential applications of cryo-EM in structural biology and rational drug discovery.

## Method

We developed a multi-modal deep learning model, called Cryo2Struct2, that integrates embeddings from the Protein Language Model (ESM) [15] with cryo-EM density maps. Cryo2Struct2 takes a 3D cryo-EM density map along with its associated amino acid sequence as inputs. To incorporate sequence information, we used the Evolutionary Scale Modeling (ESM) model [15], a pretrained language model with 3 billion parameters, to generate sequence embeddings of size 2560. These embeddings were then incorporated into the density maps for training, validation and inference.

In our previous work, Cryo2Struct [9], we trained two separate models to predict atom types and amino acid types from cryo-EM density maps. In this work, for Cryo2Struct2, we designed a unified model with a shared transformer encoder to extract features from the density map, followed by task-specific decoders for atom and amino acid type predictions. The models were trained for volumetric segmentation tasks.

The predicted probabilities for atom and amino acid types were then used to construct a Hidden Markov Model (HMM) [16], which was processed by a modified Viterbi algorithm, as introduced in Cryo2Struct [9], to align predicted voxels and build a 3D atomic protein backbone structure. We generated two atomic structures using this approach by adjusting parameters, particularly in selecting and clustering predicted C*α* atoms. These atomic structures serve as template structures for AlphaFold3, encouraging it to predict protein structures that align with the cryo-EM density map while leveraging the state-of-the art prediction capabilities of AlphaFold3.

Our approach aims to integrate cryo-EM-based structure modeling with sequence-based structure prediction, employing a late fusion strategy where the structural models build from cryo-EM is incorporated as a template for AlphaFold3. This allows AlphaFold3 to refine the structure while maintaining consistency with experimental density data, ultimately improving the accuracy of atomic model predictions.

### Model Architecture

The deep learning model is based on the 3D SegFormer [17] based architecture designed to integrate cryo-EM density map data with ESM embeddings for both amino-acid and atom-type prediction. The architecture consists of an encoder-decoder blocks, where the encoder extracts hierarchical representations of the cryo-EM density map, and the task-specific decoder separately predicts atom types and amino acid types in each voxel of cryo-EM density map.

The architecture of our deep learning module is shown in Figure 1. The encoder of the model consists of multiple Transformer [18] blocks designed to process volumetric cryo-EM density map. To fit the cryo-EM data within the memory constraints of commodity GPUs, the full 3D cryo-EM density map is divided into sub-cubes of size 32× 32 × 32. The input density map, represented as a single-channel tensor of size 32 × 32 × 32 × 1, where the last dimension corresponds to the electron density value for each voxel in cryo-EM density map, is processed through a series of convolution layers with varying kernel sizes and strides to generate multi-scale feature representations. We utilize the feed-forward multi layer perception (MLP) layer to transform the sequence embeddings generated from Protein Language Model (ESM) from 2560 to the embedding dimension of the multi-scale feature representations and add those features to the multi-scale feature representations. This helps us to integrate the ESM embeddings with the features extracted from density map, ensuring that the sequence-level information is added in the model.

**Figure 1:**
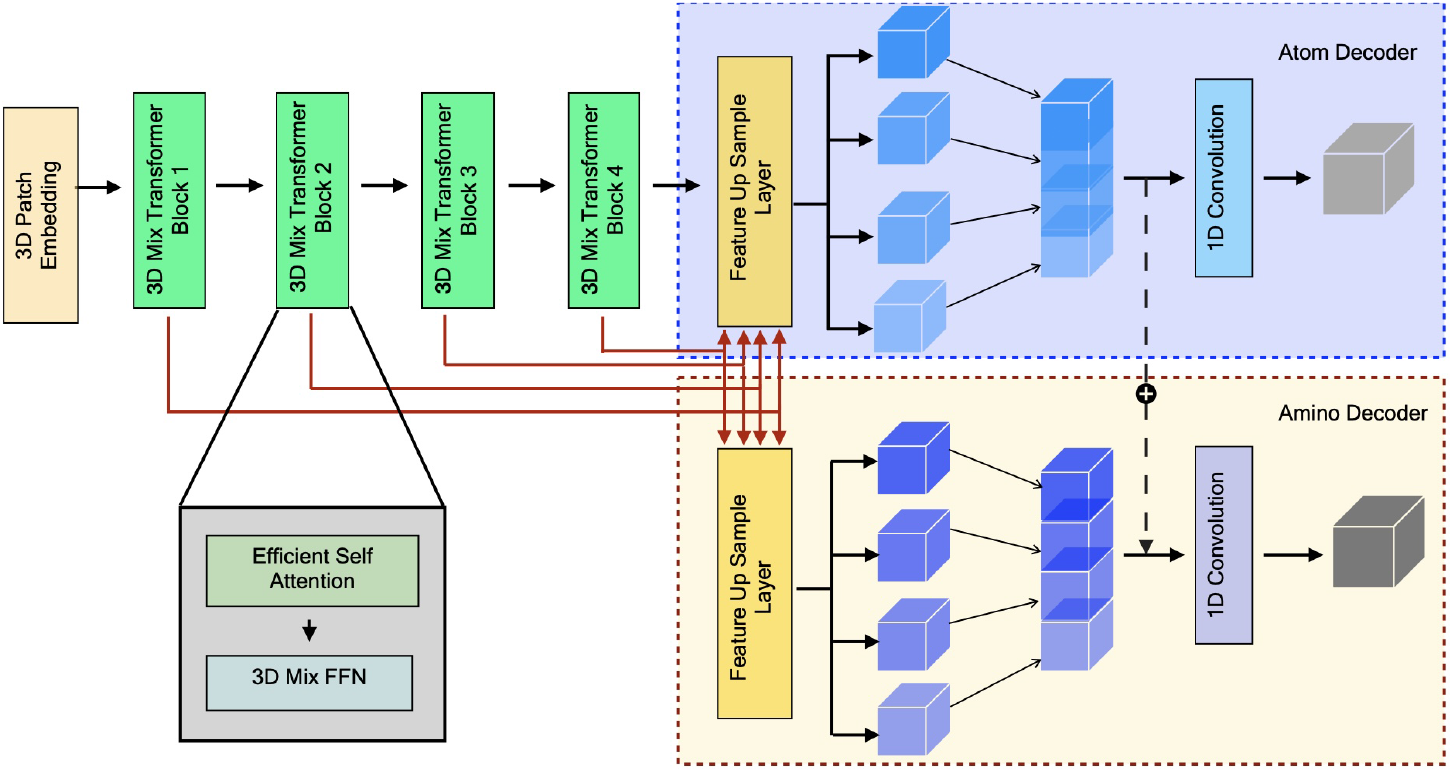
Model Overview: The architecture processes a sub-cube of a 3D cryo-EM density map using 3D Mix Transformer blocks to extract hierarchical spatial features. The Atom Decoder (blue background) classifies voxels into four atomic categories, while the Amino Decoder (yellow background) classifies voxels into 21 classes for amino acid prediction, using features from atom decoder.

Each transformer block utilizes an efficient self-attention mechanism to efficiently capturing long-range patterns in the cryo-EM density maps. The standard self-attention mechanism [18] computes attention weights and has a computational complexity of *O*(*n*^2^), however with the efficient self-attention mechanism introduced in [19, 20], the computational complexity is reduced from *O*(*n*^2^) to *O*(*n*^2^*/r*). The reduction parameter *r* is set as 4,2,1,1 in the four stages of the encoder. The Transformer bloack also contains a 3D Mix FFN as described in [19] which allows for the automatic learning of positional cues and eliminates the need for fixed encoding for better performance. Finally, the encoder outputs four feature maps, (c_1_, c_2_, c_3_, c_4_) which are used as an inputs to the decoders.

We utilize two separate decoder blocks in the model, each designed for a specific prediction task, atom-type decoder and amino acid-type decoder. The atom-type decoder processes the features (c_1_, c_2_, c_3_, c_4_) from the encoder to predict atom types into four different classes, namely, C*α*, N, C, and no presence of atoms. The amino acid-type decoder predicts amino acid types into 21 different classes by utilizing the same set of encoded features (c_1_, c_2_, c_3_, c_4_). The atom-type decoder directly predicts atomic labels from the encoded features, where as the amino acid type decoder benefits from additional features. Specifically, the amino acid type decoder incorporates the atom-type features generated from the atom-type decoder as an auxiliary feature, allowing the model to use atom-type information to improve amino acid classification accuracy.

Both atom and amino acid-type decoder follow a similar architecture, they first transform the multi-scale feature maps (c_1_, c_2_, c_3_, c_4_) from the encoder using MLP layers that project each feature layer into common embedding space. The transformed feature maps are then upsampled to a common spatial size before being fused through a convolutional layer. This fused representation undergoes dropout regularization, followed by a final convolutional layer that generates the class probability maps.

Finally, the predicted outputs from both decoders are upsampled by a factor of 4 using trilinear interpolation to get the original input size of 32 × 32 × 32 × C, where C represents output channels i.e., 4 for the backbone atom type classification (C*α*, N, C, and the absence of an atom) and 21 for the amino acid type classification (20 standard amino acids and no/unknown amino acid).

### Training and Validation

We utilized the Cryo2StructData [21] dataset which includes cryo-EM density map with the resolution in the range of [1.0-4.0 Å], to train and validate the deep learning model. The cryo-EM density maps in the dataset were released till 27 March 2023. The dataset is split according to a 90% and 10% ratio into training and validation datasets. The training dataset and validation dataset has 6652, and 740 cryo-EM density maps, respectively. Cryo2StructData [21] also provides the label maps for the cryo-EM density maps where every voxels in the density maps are labeled for both atom-type and amino acid types.

The model was trained on 32 × 32 × 32 sub-grids extracted from full cryo-EM density maps, considering only sub-grids containing at least one nonzero voxel. Training was performed using a batch size of 1000 and optimized with the Adam optimizer, which integrates parameters from the shared encoder and both task-specific decoders. We employed a weighted cross-entropy loss function (Equation 1) to address class imbalance. The model was trained using the Distributed Data Parallel technique across four GPUs, each with 80 GB of memory, and a learning rate 10^*−*4^ [22].

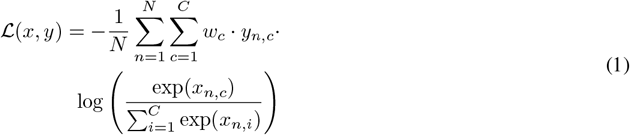

where, ℒ (*x, y*) represents the weighted cross-entropy loss. *N* is the number of samples in the minibatch. *C* is the number of classes. *w*_*c*_ is the weight for class *c* computed using Formula 2. *x*_*n,c*_ is the logit for class *c* in sample *n*, and *y*_*n,c*_ is a binary indicator (0 or 1) of whether class *c* is the correct classification for sample *n. ω*_*c*_ in Formula 2 represents the weight assigned to class *c, n*_*c*_ is the number of samples in class *c*, and 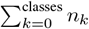 is the total number of samples across all classes. The model with the lowest validation loss is selected as the trained model for inference stage.

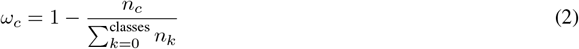

### Clustering predicted C*α* voxels

The deep learning model predicts C*α* atoms from cryo-EM density maps. Given the challenges of this prediction task, it is common for multiple spatially close voxels to be predicted as individual C*α* atoms. To address this redundancy, we performed clustering of the predicted C*α* atoms based on their proximity. Specifically, we applied two clustering thresholds: 2 Å and 3 Å. The 3 Å clustering threshold is more rigid, as the average distance between two C*α* atoms is approximately 3.8 Å. By using a 3 Å threshold, the clustering helps to group voxels that are within 3 Å of each other, which assists in improving alignment and reducing incorrect connections between C*α* atoms. The co-ordinate of the clustered C*α* in each cluster, and the probabilities of the C*α* amino acid-type for each cluster are averaged and are used for constructing the Hidden Markov Model (HMM).

### HMM and Customized Viterbi

We utilized the same approach used by Cryo2Struct [9] to align the protein amino acid sequence to the predicted and clustered C*α* voxel co-ordinates. The transition probability for the HMM is constructed based on the distance between two predicted C*α* voxels in the 3D space, calculated using the Euclidean distance formula. The distance is converted into a probability using the modified Gaussian probability density function (PDF) with a mean (*µ*) of 3.8047 Å and a standard deviation (*σ*) of 0.036 Å. Both *µ* and *σ* were estimated from the distances between two adjacent C*α* atoms in the known protein structures in the training dataset. Additionally, we introduce a fine-tune able scaling factor (Λ) that multiplies with (*σ*) to make the model adjustable. We set (Λ) to 10.

The emission probability matrix (*δ*) for each C*α* state (voxel) is calculated from both its predicted amino acid type probability and the background (prior) probability of 20 amino acids in the nature. Specifically, the geometric mean of the two is calculated as 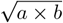, where *a* corresponds to the predicted probability for each amino acid type, and *b* represents the background frequency of the amino acid type, that was precomputed from the true protein structures in the training dataset. The geometric means for 20 amino acid types are normalized by their sum as their final emission probability.

The initial probability for a C*α* state is the probability that it generates the first amino acid of the protein sequence normalized by the sum of these probabilities of all the C*α* states.

The customized Viterbi algorithm is used to find the most likely path in the HMM to generate a protein sequence with the maximum probability. For a multi-chain protein complex, the sequence of each chain is aligned with the HMM one by one. Once a chain is aligned, the states in the hidden path aligned with it are removed from the HMM before another chain is aligned. In the alignment process, it is ensured that any C*α* state occurs at most once in one hidden state path. One distinct strength of this HMM-based alignment approach is that every amino acid of the protein is assigned to a C*α* position as long as the number of the predicted C*α* voxels is greater than or equal to the number of the amino acids of the protein.

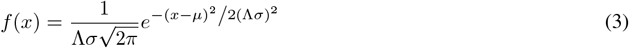

### Templates for AlphaFold3

To improve the accuracy of structure prediction, we integrated template-based modeling into AlphaFold3 using Cryo2Struct2-generated structures as templates. Our approach involves aligning query protein sequences with template sequences derived from cryo-EM-based structural predictions, in this case Cryo2Struct2. The alignment process extracts corresponding residue indices between the query and template sequences, ensuring a structurally meaningful template mapping. The extracted templates are then formatted into a JSON-compatible file for AlphaFold3, which includes multi-chain sequence entries, alignment indices, and embedded mmCIF template data. By incorporating these templates, AlphaFold3 can leverage prior structural information, improving alignment with experimentally determined structures.

## Results

The test dataset for atomic structure prediction was collected from the Electron Microscopy Data Bank (EMDB) [23] and consists of cryo-EM density maps deposited in the year 2024. These test data were not included in the training of either AlphaFold3 [14] or Cryo2StructData [21], ensuring an unbiased evaluation of model performance. The training data cut-off dates for AlphaFold3 and Cryo2StructData were September 2021 and March 2023, respectively.

The test dataset consists of 61 cryo-EM density maps with an average resolution of 3.10 Å, ranging from 2.4 Å to 3.9 Å. The average number of residues per structure is 1,230.8, with a range of 374 to 3,245 residues. To assess structural similarity, we used the standard TM-score, which quantifies how well a predicted model aligns with its corresponding known structure. TM-scores were calculated using US-align [24], a protein complex structure comparison tool, with options enabled for aligning multi-chain oligomeric structures and all chains, as recommended for biological assembly alignment.

To ensure a fair comparison between models of varying lengths, the global TM-score was normalized using the length of the corresponding experimental structure. The TM-score ranges from 0 to 1, with 1 representing a exact structure match.

In terms of structural accuracy, as shown in Figure 2 the average TM-score for AlphaFold3 using Cryo2Struct2 multi-template guidance is 0.32, whereas AlphaFold3 without template guidance achieves an average TM-score of 0.28.

**Figure 2:**
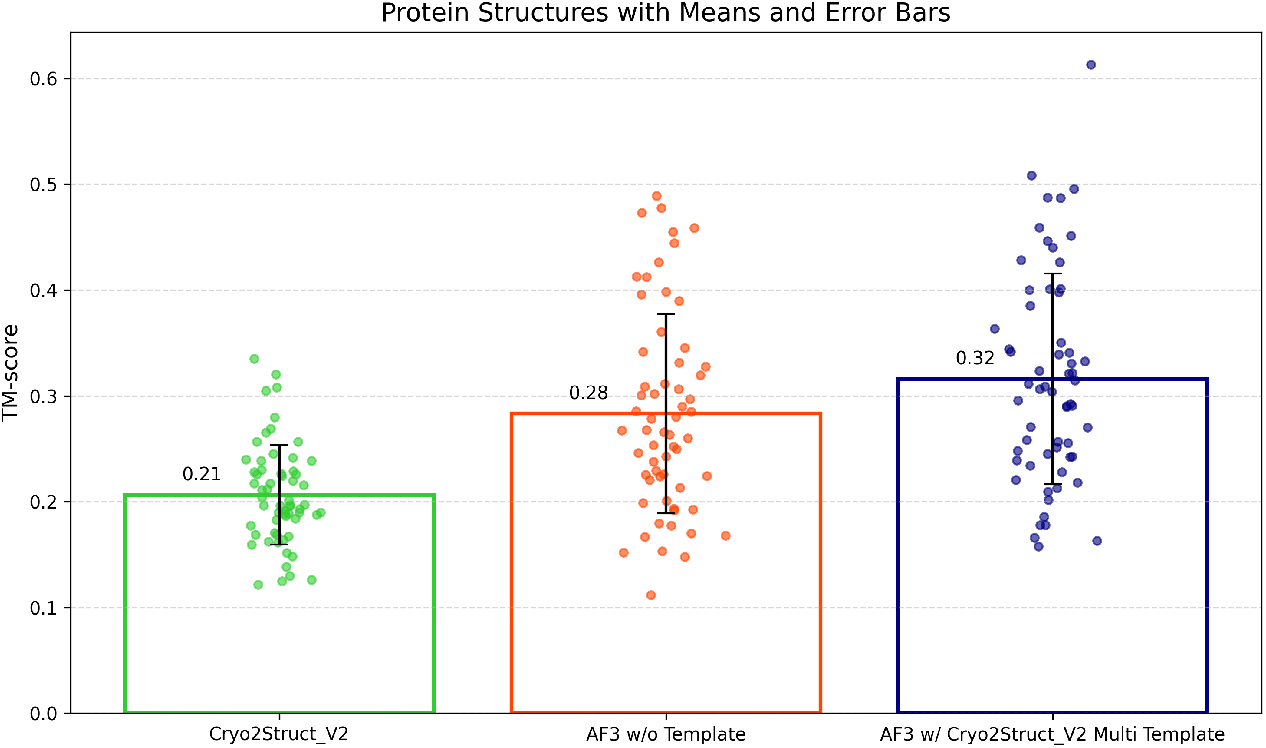
Evaluation of 61 protein structures modeled by each method using TM-Score. Each dot represents a structure predicted by the respective approach. The mean scores for structures modeled by Cryo2Struct2, AlphaFold3 without a template, and AlphaFold3 with multiple Cryo2Struct2 templates are 0.21, 0.28, and 0.32, respectively.

The average TM-score for structures modeled directly from Cryo2Struct2 alone is 0.21. These results indicate that integrating Cryo2Struct2-generated templates with AlphaFold3 improves structural accuracy, highlighting the benefit of incorporating cryo-EM-derived information for refining atomic structure predictions.

In terms of sequence accuracy, as shown in Figure 3, sequence identity is measured as the proportion of identical residues among the aligned residues. The average sequence identity (sequence-ID) for AlphaFold3 using Cryo2Struct2 multi-template guidance is 0.35, whereas AlphaFold3 without template guidance achieves an average sequence-ID of 0.25. For structures modeled directly from Cryo2Struct2 alone, the average sequence-ID is 0.12. These results indicate that incorporating Cryo2Struct2 templates into AlphaFold3 enhances sequence accuracy, further supporting the effectiveness of template-based refinement in cryo-EM-guided protein structure modeling.

**Figure 3:**
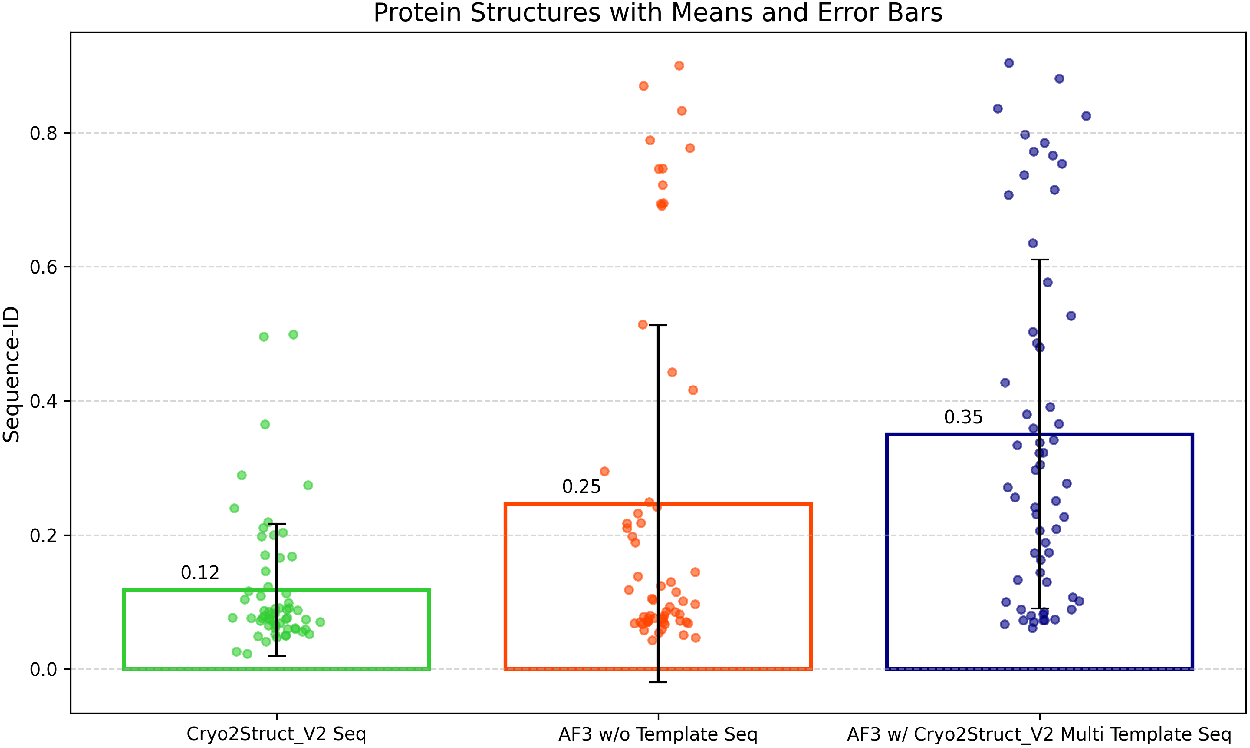
Evaluation of 61 protein structures modeled by each method using Sequence identity. Each dot represents a structure predicted by the respective approach. The mean scores for structures modeled by Cryo2Struct2, AlphaFold3 without a template, and AlphaFold3 with multiple Cryo2Struct2 templates are 0.12, 0.25, and 0.35, respectively.

Figures 4, 5, and 6 showcase examples of modeled structures, providing detailed insights into their folds. In Figure 4, the structure predicted by AlphaFold3 without template guidance shows a lower match with the experimentally determined structure, with a TM-score of only 0.265. In contrast, when guided by Cryo2Struct2, the predicted structure aligns significantly better with the PDB-deposited structure, resulting in a higher TM-score of 0.398.

**Figure 4:**
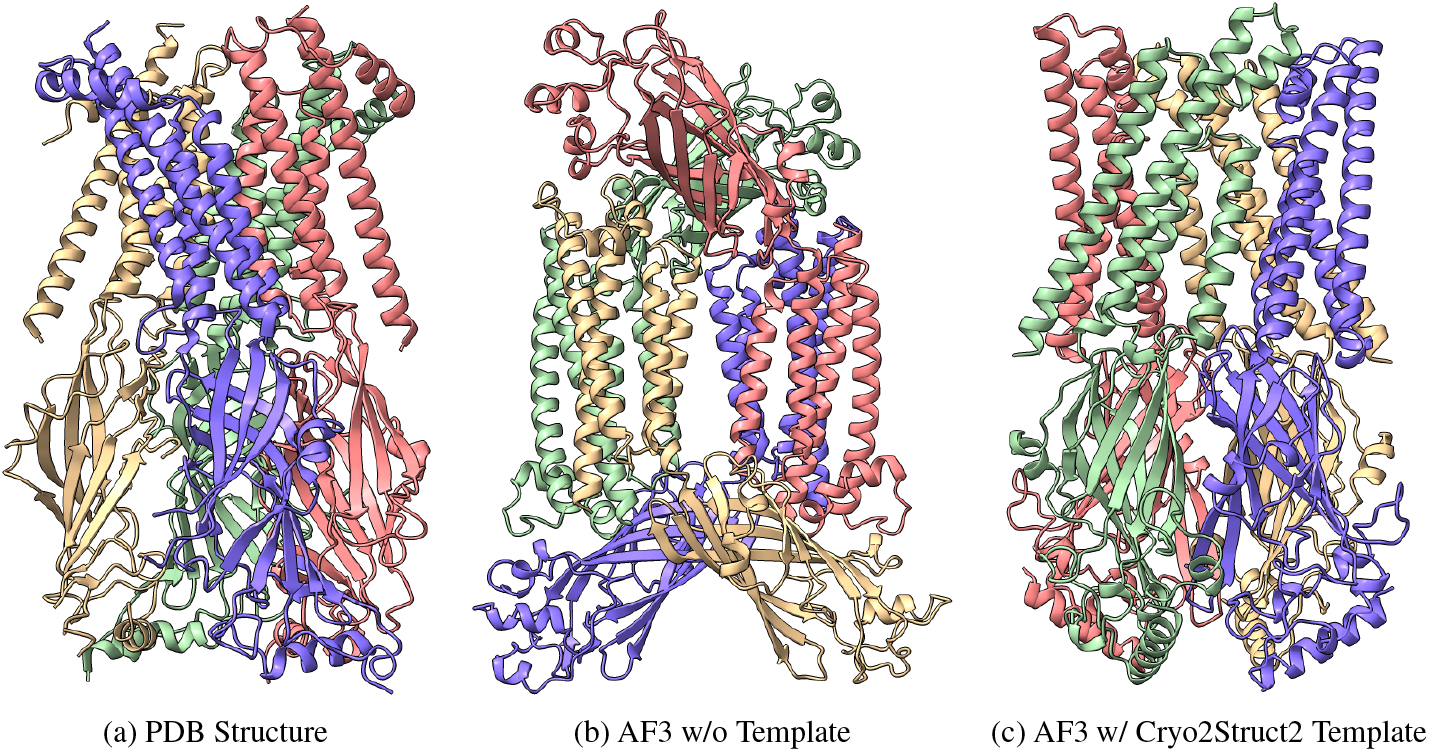
Example of modeled structures. **(a)** PDB-deposited structure (PDB Code: 8C1W). **(b)** AlphaFold3 prediction without templates has TM-Score of 0.265. **(c)** AlphaFold3 prediction using a Cryo2Struct2-generated template has TM-Score of 0.398.

**Figure 5:**
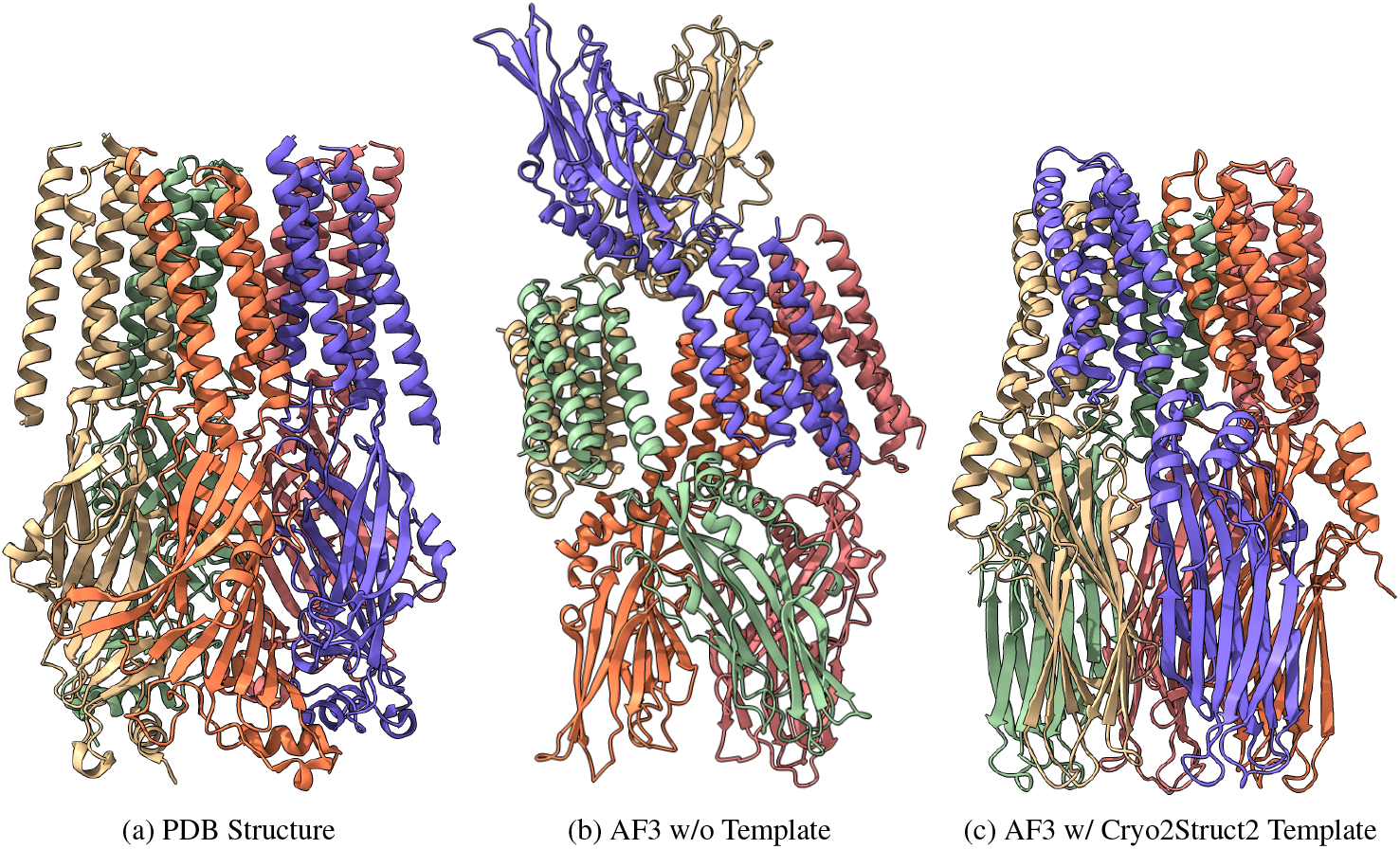
Example of modeled structures. **(a)** PDB-deposited structure (PDB Code: 8BL8). **(b)** AlphaFold3 prediction without templates has TM-Score of 0.278. **(c)** AlphaFold3 prediction using a Cryo2Struct2-generated template has TM-Score of 0.400.

**Figure 6:**
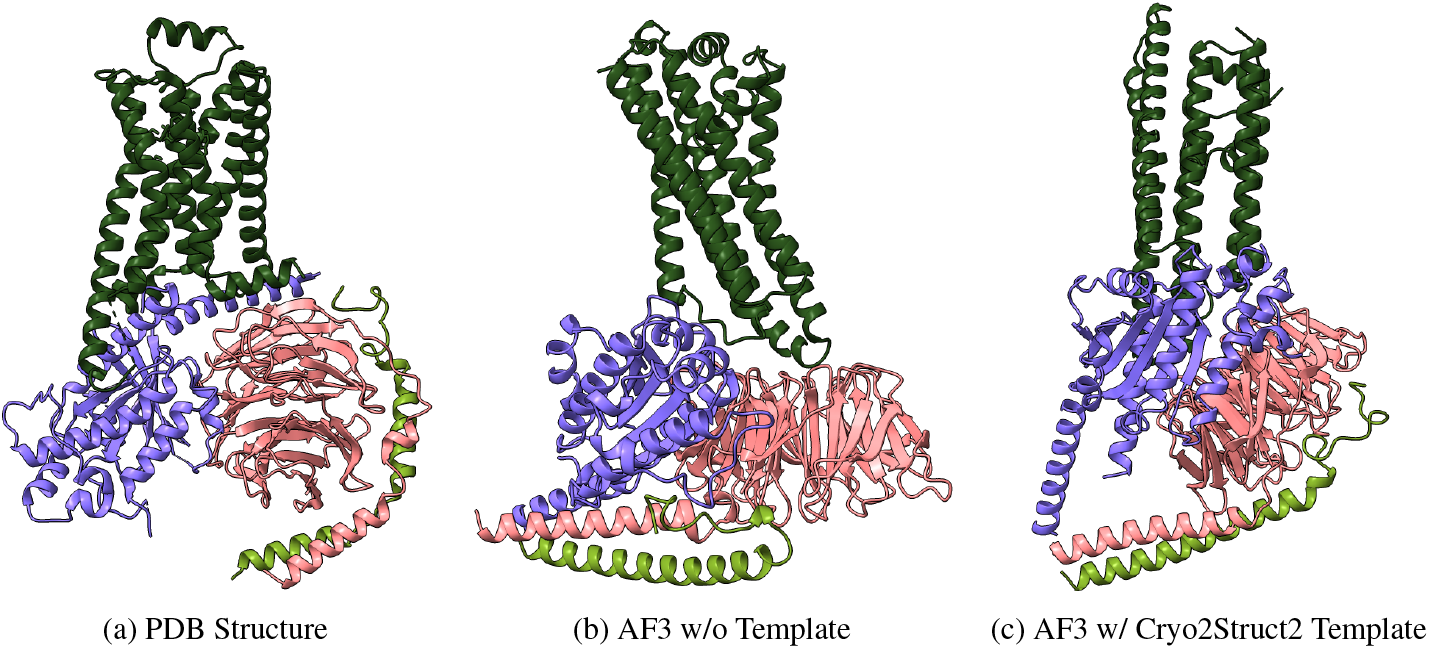
Example of modeled structures. **(a)** PDB-deposited structure (PDB Code: 8GE1). **(b)** AlphaFold3 prediction without templates has TM-Score of 0.412. **(c)** AlphaFold3 prediction using a Cryo2Struct2-generated template has TM-Score of 0.451.

A similar trend is observed in Figure 5, where the AlphaFold3 prediction without template guidance yields a TM-score of 0.278, indicating a lower structure match. However, the structure predicted using Cryo2Struct2 guidance demonstrates improved alignment with the reference structure, achieving a higher TM-score of 0.4. These examples highlight the effectiveness of integrating Cryo2Struct2 templates in correcting and improving structure accuracy.

## Discussions

In this work, we demonstrated the effectiveness of using cryo-EM-based predicted atomic structures as templates for AlphaFold3 to refine and correct protein structures, ensuring they accurately represent the atomic details derived from the cryo-EM density maps. By leveraging only the templates generated from cryo-EM predictions, we were able to predict structures that closely correspond to the experimental density maps, which holds great potential for identifying novel proteins and understanding biological mechanisms relevant to drug discovery. AlphaFold3’s ability to refine structures is particularly valuable in regions where the cryo-EM-based method struggles, such as areas with low resolution and missing electron density values that typically complicate accurate structure modeling.

In the future, the approach outlined here can be extended to support the prediction of joint structures of complexes, including proteins, nucleic acids, small molecules, ions, and modified residues. By integrating cryo-EM data with

AlphaFold3’s advanced structure modeling capabilities, we have developed a versatile framework that enables the generation of complex structures, broadening the scope of cryo-EM-based complex structure modeling.

Our experiments highlight that AlphaFold3 not only refines cryo-EM-based structures but also benefits from these cryo-EM predictions as templates, creating a mutually reinforcing feedback loop that improves both methods. Additionally, the alignment strategy implemented in Cryo2Struct [9] has proven to be effective as a standalone tool, with potential for adaptation to other deep learning models. Future work will focus on further improving this alignment strategy to improve the quality and accuracy of generated atomic structural models.

## Code availability

The source code for Cryo2Struct2 is available in the GitHub repository: https://github.com/BioinfoMachineLearning/Cryo2Strut2.git.

## Acknowledgements

This work was supported in part by an NIH grant (R01GM146340) to JC. The computation for this work was performed on the high performance computing infrastructure provided by Research Computing Support Services at the University of Missouri, Columbia MO.

